# Visual stimulation with blue wavelength light drives V1 effectively eliminating stray light contamination during two-photon calcium imaging

**DOI:** 10.1101/2021.02.27.433182

**Authors:** Tatiana Kuznetsova, Kamil Antos, Evgenya Malinina, Stylianos Papaioannou, Paolo Medini

**Author notes:** equal contribution (Tatiana Kuznetsova, Kamil Antos). **Correspondence:**, hus H, Johan Bures väg 12, Biologihuset, Umeå universitet, 901 87 Umeå.

## Abstract

**BACKGROUND:** Brain visual circuits are often studied *in vivo* by imaging Ca^2+^ indicators with green-shifted emission spectra. Polychromatic white visual stimuli have a spectrum that partially overlaps indicators’ emission spectra, resulting in significant contamination of calcium signals.

**NEW METHOD:** To overcome light contamination problems we choose blue visual stimuli, having a spectral composition not overlapping with Ca^2+^ indicator’s emission spectrum. To compare visual responsiveness to blue and white stimuli we used electrophysiology (visual evoked potentials–VEPs) and 3D acousto-optic two-photon(2P) population Ca^2+^ imaging in mouse primary visual cortex (V1).

**RESULTS:** VEPs in response to blue and white stimuli had comparable peak amplitudes and latencies. Ca^2+^ imaging revealed that the populations of neurons responding to blue and white stimuli were largely overlapping, that their responses had similar amplitudes, and that functional response properties such as orientation and direction selectivities were also comparable.

**COMPARISON WITH EXISTING METHODS:** Masking or shielding the microscope are often used to minimize the contamination of Ca^2+^ signal by white light, but they are time consuming, bulky and thus can limit experimental design, particularly in the more and more frequently used awake set-up. Blue stimuli not interfering with imaging allow to omit shielding without affecting V1 physiological responsiveness.

**CONCLUSIONS:** Our results show that the selected blue light stimuli evoke physiological responses comparable to those evoked by white stimuli in mouse V1. This will make complex designs of imaging experiments in behavioral set-ups easier, and facilitate the combination of Ca^2+^ imaging with electrophysiology and optogenetics.

**Highlights:**

- White and blue light trigger VEPs with similar amplitudes and latencies in mouse V1
- Blue-and white-responding neurons are two largely overlapping neuronal populations
- Blue and white evoke Ca^2+^ responses similar in magnitude and latency
- Blue and white evoke Ca^2+^ responses similar in orientation/direction selectivity
- Blue stimuli could be an alternative to white ones in behavior and opto-physiological tests

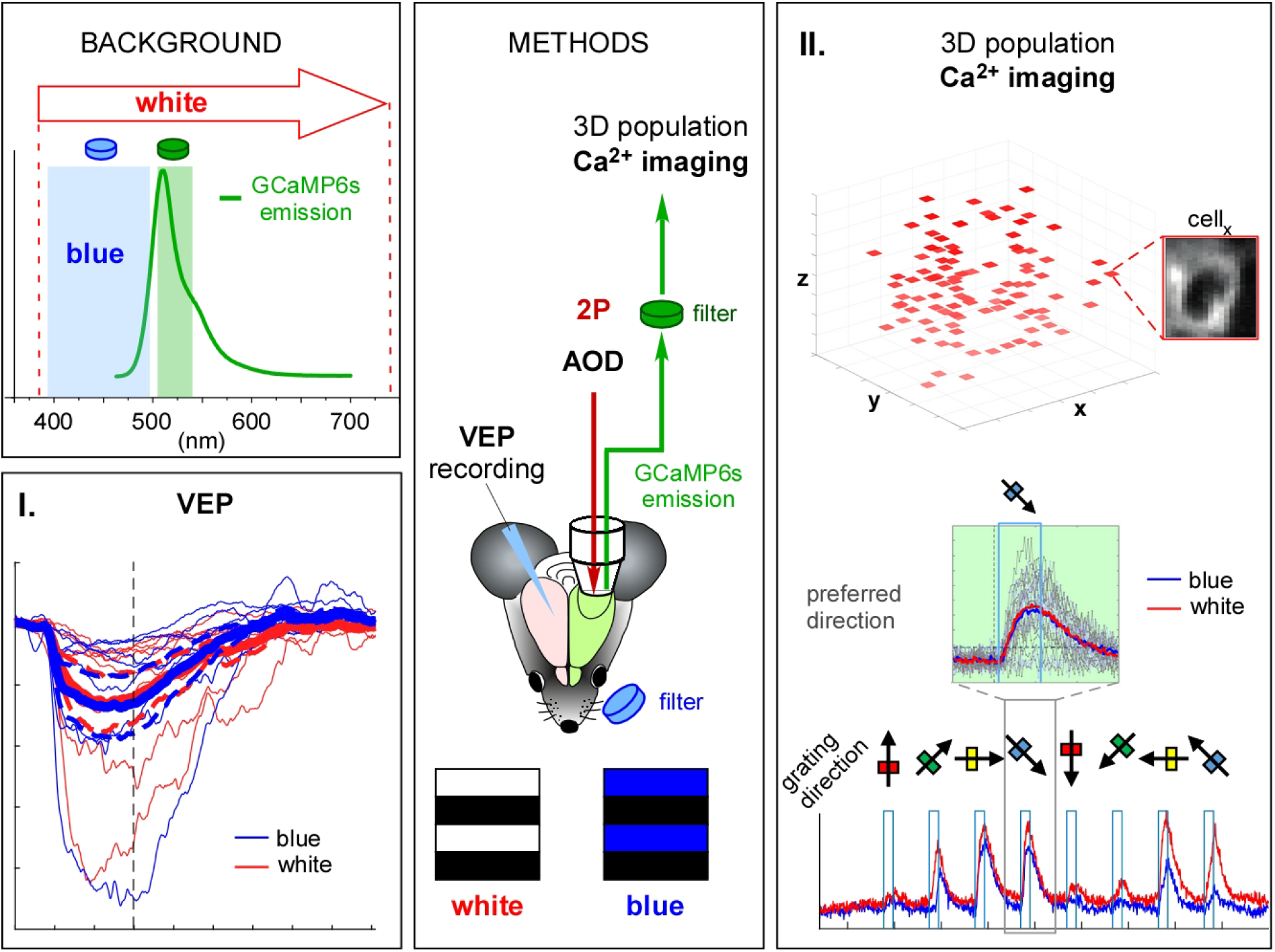

## 1. Introduction

Ca^2+^ imaging is widely used to monitor neuronal responses to different stimuli and conditions at population, axonal, dendritic and dendritic spine levels both *in vitro* and *in vivo* (Dana et al., 2019; Grienberger and Konnerth, 2012). Somatic and axonal calcium transients accompany neuron’s spike output, whereas dendritic Ca^2+^ signals can identify local and global dendritic activations driven by synaptic input as well as by back-propagating action potentials (Ali and Kwan, 2019). Combination of Ca^2+^ imaging with optogenetic tools permits to manipulate the activity of specific neuronal populations with high spatiotemporal precision, allowing to dissect the functional organization of neural circuits in different model organisms (e.g. flies, worms, fish and mammals, spanning from mice to non-human primates) and in various brain regions (Carrillo-Reid et al., 2017; Galvan et al., 2017; Jiao et al., 2018; Shipley et al., 2014; Simpson and Looger, 2018).

Changes in intracellular Ca^2+^ concentration are estimated by changes in fluorescence of Ca^2+^ indicators, emitting the light of certain wavelengths if activated by excitation light when bound to calcium. Currently genetically-encoded calcium indicators (GECIs) are the most used, as they can also be stably expressed in transgenic animals (Dana et al., 2014; Dana et al., 2018). The most commonly used GECIs GCaMPs – have calmodulin as a Ca^2+^ sensor and cpEGFP (circularly permuted enhanced green fluorescent protein) as a reporter. cpEGFP has a green emission spectrum peaking at around 515 nm, when GCaMP is bound to Ca^2+^. Recently developed red-shifted GECI – such as jRGECO1a, jRCaMP1a and b have emission spectra peaking at around 600 nm (Dana et al., 2016; Dana et al., 2018).

Regardless of GECI used, the detection of activity-driven changes in its fluorescence can be hindered by light contamination (Leinweber et al., 2014). Indeed, the spectra of ambient light (e.g. from monitor screens) overlaps with the emission spectra of GECIs. For example, the white light of CRT (cathode-ray tube) screens, often used for visual stimuli presentation, is formed by a combination of red, green, and blue primary colors, spreading between 380 and 730 nm (Fig. 1 A). Light contamination is particularly problematic in the case of visual stimulation, used to explore for example the visual cortex (both primary and associative), a classical model to investigate the function and plasticity of cortical microcircuits *in vivo* (Yang et al., 2018). In addition, visual stimuli are used in awake set ups as visual cues (Goard et al., 2016; Hwang et al., 2019), in visually guided eye movement tasks (Itokazu et al., 2018) and in virtual reality systems when the functions of (pre)motor (Leinweber et al., 2017) and associative (Krumin et al., 2018) cortical networks are investigated.

**Figure 1.**
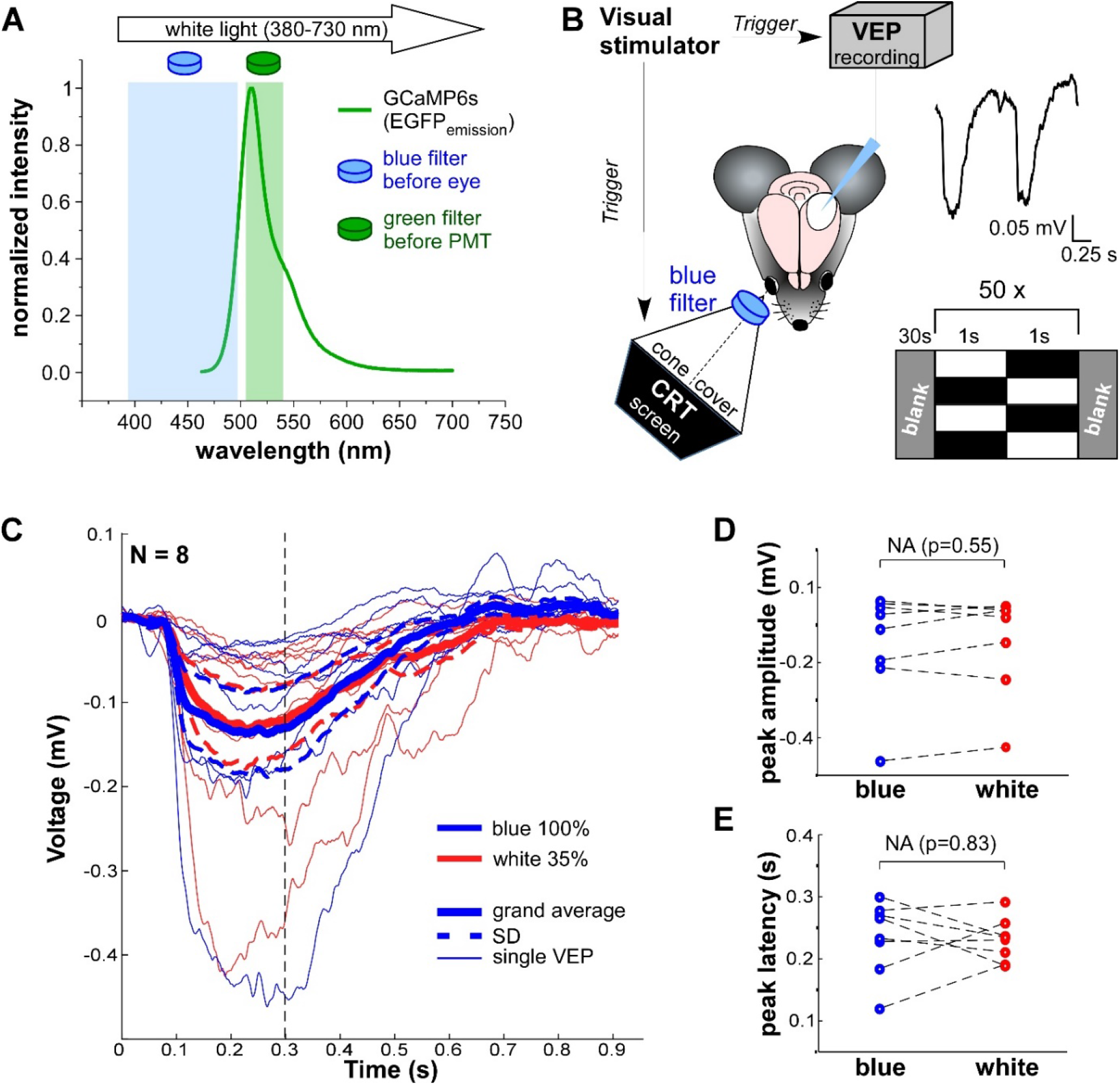
Blue/black gratings evoke visual evoked potentials with similar amplitudes and peak latencies as white/black ones. **A**. Emission spectra of GCaMP6s overlaid with spectra of light passing through the blue filter (FESH0500, 392-497 nm), through the green filter (FBH520-40, 503-539 nm) located before PMT for acquisition of GCaMP6s fluorescence during Ca^2+^ imaging and with spectra of white light (380-730 nm) coming from the CRT screen. **B**. Sketch of VEP experiment. On the left: position of mice eye in relation to CRT screen, covered by opaque cover. Blue filter is inserted in the slit at the tip of the cover. VEPs are recorded in the contralateral to the eye V1. In *italic*: triggering of HEKA by Visual stimulator (ViSaGe). On the right, below: scheme of visual stimulation protocol appearing on the CRT screen. It consists of 50 repetitions of stationary gratings alternating in counter-phase (every 1s), equiluminant to the gratings gray is presented 30 s before and after 50 repeats; on the right, above: representative VEP (averaged across the repeats), temporally aligned to the presented gratings, shown below. **C**. Grand averages (N=8, thick lines) ± SD (thinner dashed lines) and individual VEPs (for single animals; thin lines) in response equi-powerful blue/black (100% intensity; in blue) and white/black (100% intensity; in read) gratings. Vertical black dashed line confines the time for peak search (0.3 s). Note that the responses are strongly overlapping. **D, E**. Amplitude (D) and latency (E) of single VEP’s peaks in response to blue/black (in blue) and white/black (in read) gratings. Notice, that there is no statistical significant difference in response to blue/black and white/black gratings neither in amplitude, nor in latency (p=0.55, Wilcoxon signed rank test, and p=0.83, paired t-test 0.84, respectively; N=8).

A common way to bypass this problem is to mask the gap between objective and imaged brain area with opaque material (Leinweber et al., 2014). This limits the experiment to an optical approach, excluding (or rendering problematic) simultaneous electrophysiological recording, optogenetic manipulation of an area outside the imaged one, or drug application. Alternatively, one can shield the objective by placing black curtains (often close to the preparation) between mice eye and craniotomy. This solution however might be bulky or mechanically undoable, preventing or creating problems in open frame behavioral experiments, and sometimes might not be sufficient.

Another approach to overcome the light contamination is the synchronization of the screen’s or projector’s (that are generating the visual signal) output to the turnaround time of the laser scanning electronics (when the signal is not collected (Leinweber et al., 2014; Reiff et al., 2010)). This elegant solution requires engineering for desirable synchronization for each setup and restricts to stick to certain scanning speed (not suitable for all imaging systems).

The third approach, considered here, is to substitute the white light (used for visual stimuli and causing the light contamination) with light that has the spectrum spectrally separated from GECI emission. Taking into account the current existence of “green” and “red” GECI, blue light (~395/400-495/500 nm) could be a good option (Fig 1A).

However, although visual stimulation with blue light (400-500 nm) has been used during in vivo imaging (Lecoq et al., 2014), it has not been systematically studied whether visual stimulation with blue light, selected to minimize the contamination of the recorded “green” calcium indicator signal (500-540 nm), evokes visual responses similar to the ones triggered by the white light in primary visual cortex (V1) of experimental rodents. Rodents, in particularly mice, are primarily nocturnal animals with dichromatic color vision and rod-dominated retinas (Huberman and Niell, 2011). Here we tested whether visual responsiveness and functional response properties of visual V1 neurons (e.g. orientation and directional selectivity) are similar in response to white and blue stimuli. It is possible that the blue light could trigger a weaker response, since the wavelength composition of blue light covers a smaller part of mice spectral sensitivity function, as obtained from electroretinogram (ERG) and behavioral tests (Jacobs et al., 1991; Jacobs et al., 2004; Rocha et al., 2016). On another hand, blue and white lights should activate two main pigments with similar absorption spectra in retinal photoreceptors: rhodopsin in rods and the “green” long/middle (L/M) wave-sensitive opsin in cones (but very weakly shortwave/ultraviolet (S/UV)-sensitive opsin), as predicted by their spectral sensitivity functions (Peirson et al., 2018).

To address this question, we recorded VEPs and performed 3D acousto-optic 2P population Ca^2+^ imaging (that allows to monitor the activity of chosen multiple individual neurons) in V1 of mice during the presentation of white/black and blue/black gratings, and estimated if the responses to this two color-types of gratings are significantly differed from each other.

## 2. Materials and methods

### 2.1. Animal strain and housing

For LFP experiments young adult (1-2 months) C57BL/6J male wild type mice were used. For 2-photon Ca^2+^ imaging of neuronal population young adult (1-3 months) Thy1-GCaMP6s mice (C57BL/6J-Tg(Thy1-GCaMP6s)GP4.3Dkim/J; The Jackson Laboratory, N 024275) were used (Dana et al., 2014). Mice were housed in Umeå Center for Comparative Biology’s (UCCB) and maintained in a 12 h light/dark cycle, with food and water available *ad libitum*. At the end of the experiment animals were sacrificed by cervical dislocation. All procedures were done according to the Ethical permit from the Swedish Ethical Committee for Northern Sweden.

### 2.2. In-vivo extracellular electrophysiology

#### 2.2.1. Mouse preparation and surgery

Animals were anesthetized with isoflurane (4% for induction; 1.5-2.5 during surgery). To reduce the inflammatory response and brain edema Dexamethasone (0.7 mg/kg, *i*.*m*.; 60-100 µl of 0.2 mg/ml in saline) was administered before the surgery. To protect the cornea from dehydration, we applied artificial tears (VISCOTEARS 2 mg/g; FASS) on the eyes and covered them with Ø 6 mm cover glass. Body temperature during the experiments was constantly monitored with a rectal thermocouple probe and maintained at 37 °C with a feedback-controlled heating blanket (using ATC-2000 Animal Temperature Controller).

After removing the skin and periosteum, a metal head-plate was glued to the bone with Superglue in the hemisphere opposite to the planned craniotomy. Then a recording bath from dental cement (Paladur) was formed at the borders of the skull. The attachment of the head-plate and the bath was reinforced by tissue glue (3M Vetbond). The bath allowed to keep the cortical surface always covered by physiological solution (0.9% NaCl) and to place the ground electrode. After cement polymerization the bone above the V1 was carefully thinned by drilling and a small rectangular craniotomy was cut with a surgical blade ca 3 mm mediolateral from lambda (above the binocular portion of V1).

#### 2.2.2. Local field potentials (LFP) recordings

A glass micropipette (1-2 MΩ; Ø ca 10 µm) filled with saline (0.9% NaCl) was positioned at a cortical depth of 200 µm. LFP were recorded using an ELC-03XS amplifier (NPI Electronic; Germany) and acquired via HEKA LIH 8+8 data acquisition interface and PATCHMASTER software (HEKA; Germany). Signals were amplified (x1000), bandpass-filtered (0.1–100 Hz) and digitized (1 kHz). LFP recordings were triggered by the onset of visual stimulus (reversal of the stationary gratings, see below) appearing on the CRT screen and controlled by a stimulus generator (ViSaGe from Cambridge Research Systems Ltd; UK; see Fig.1B, left). Each recording (1 sweep) lasted 1.91 s (grating pattern reversed at 0 and 1 s) and 50 such sweeps were recorded within a series (see Fig.1B, upper right).

Series with white/black and blue/black grating presentations were recorded in a random order in different animals.

#### 2.2.3. Visual stimulation during LFP recording

We presented stimuli on a calibrated video monitor system (ViSaGe MKII Stimulus Generator, Cambridge Research Systems Ltd, CRT monitor HP P1230, refresh rate 140Hz, color temperature 9300 K), controlled by custom-made scripts in Matlab. The CRT screen was light-shielded by the opaque cone with a round slit on its end for blue filter insertion. This slit was positioned in close vicinity to the mouse eye (contralateral to the craniotomy), with the CRT screen located at ca 40 cm from the eye. Visual stimuli (see Fig.1B, lower right) were stationary gratings alternating in counter-phase (horizontally oriented; with squared spatial profile, spatial frequency 0.05 cycles/degree, and temporal frequency 0.5 Hz). White/black or blue/black gratings with equal output power (ca 3 µW, measured by silicon power meter - Thorlabs PM100D, at 40 cm distance from the screen, on the level of the slit) were presented. To reach similar output power levels for these two visual stimuli we used the following configurations: When white/black gratings were used, the intensity of the white on the CRT screen was set to 35% and the filter was removed from the slit in the cone. In case of blue/black gratings intensity of white was set to 100% and a blue filter (FESH0500, Thorlabs, bandpass: 392-497 nm) was inserted into the slit.

##### LFP data processing

LFP responses to each pattern reversal within a series (100 reversals, 50 sweeps – 2 reversals/sweep) were averaged so to measure VEP responses for both color-types of gratings. The first 40 ms of each series were considered as a baseline, as there is not even retinal ganglion cell activation during this time window (Mazzaro et al., 2016), and no synaptic activation of layer 4 V1 neurons (Medini, 2011). The average voltage reading during 0-40 ms window was subtracted from each averaged trace (allowing for VEP peak search and alignment of the different series for grand average calculation). This was done in a custom-made software, written in MatLab. VEP peaks were searched within 300 ms post-stimulus onset (pattern reversal) in OriginPro 2017 software; peak amplitudes and latencies were determined. For grand average calculations, VEPs of individual animals (N=8) in response to white/black or blue/black gratings were averaged and grand mean ± standard deviation (SD) were plotted (see Fig. 1C).

### 2.3. 3D Acousto-optic 2-photon Ca^2+^ imaging of neuronal populations

#### 2.3.1. Mouse preparation and surgery

Surgical anesthesia and animal preparation were done as described for LFP experiments. Part of the skin was cut out, the periosteum was removed and a circular titanium head post was attached to the skull using superglue, tissue glue and dental cement with binocular V1 in the center (3 mm mediolateral from lambda). This head post was used later to create a bath chamber, filled with saline warmed to body temperature (for IOI) or with ultrasound gel (for Ca^2+^ imaging). In the middle of this bath, a circular craniotomy (ca 4 mm in diameter) was made. To cover the craniotomy, a round glass coverslip (Ø 5 mm) was glued to the bone with a tiny amount of superglue. Since isoflurane (1-1.5%) anesthesia can affect the patterns of Ca^2+^ activity in mouse V1 (Goltstein et al., 2015), for the recording of visually-evoked activity we switched to a lighter anesthesia regimen: with chlorprothixene (1 mg/kg, 60–100 µl of 0.3 mg/ml in saline, i.m.; (Nauhaus et al., 2012)) and 0.5% isoflurane that was gradually reduced after the surgery (Juavinett et al., 2017).

#### 2.3.2. Intrinsic optical imaging (IOI)

We used IOI to identify the location of V1 relative to the blood vessel pattern. For images acquisition an Optical Imager 3001 system (Optical Imaging Inc, Rehovot, Israel) was used, equipped with: (1) HAL 100 illuminator with quartz collector (423000-9901-000) as a light source with DC power supply (TPR3020S, Atten instruments), (2) a macroscope consisting of 2 tandem Nikon lenses (NIKKOR 50mm f/1.2 and AF DC-NIKKOR 135mm f/2D) attached to 25 Hz CCD-camera (A-1000m/D, Adimec, 237127) and (3) Vdaq software for data acquisition. The Optical imaging Vdaq software triggered the Visual stimuli generator (ViSaGe) via custom-written routine in Matlab.

A “green” vascular image was captured while illuminating the cortex with 546 ± 30 nm light (bandpass filter 91933, Thorlabs) with the pia surface in focus (see Fig. 2A, upper left). To obtain activity maps (see Fig. 2A lower left), images were acquired with monochromatic 630 nm ± 10 nm light filter (Thorlabs), defocused ca 500–600 μm below the pial surface, while the lenses diaphragm was open. Image size was approximately 2.7 x 2.7 mm and 996 x 996 pixels; 3×3 spatial online binning was applied. During 10 s of stimulation 50 data frames were acquired (200 ms each, consisting of 5 camera frames). During IOI drifting gratings of 0 or 90 degree (sine waveform, spatial frequency 0.03 cycles/degree, drift velocity 1 cycle/sec; 100% contrast) or equi-luminant blanks were displayed on the screen. Stimuli (0/90 degree grating and the blank) were presented in random order with 15s of inter-stimulus interval. The protocol was repeated 10 times.

**Figure 2.**
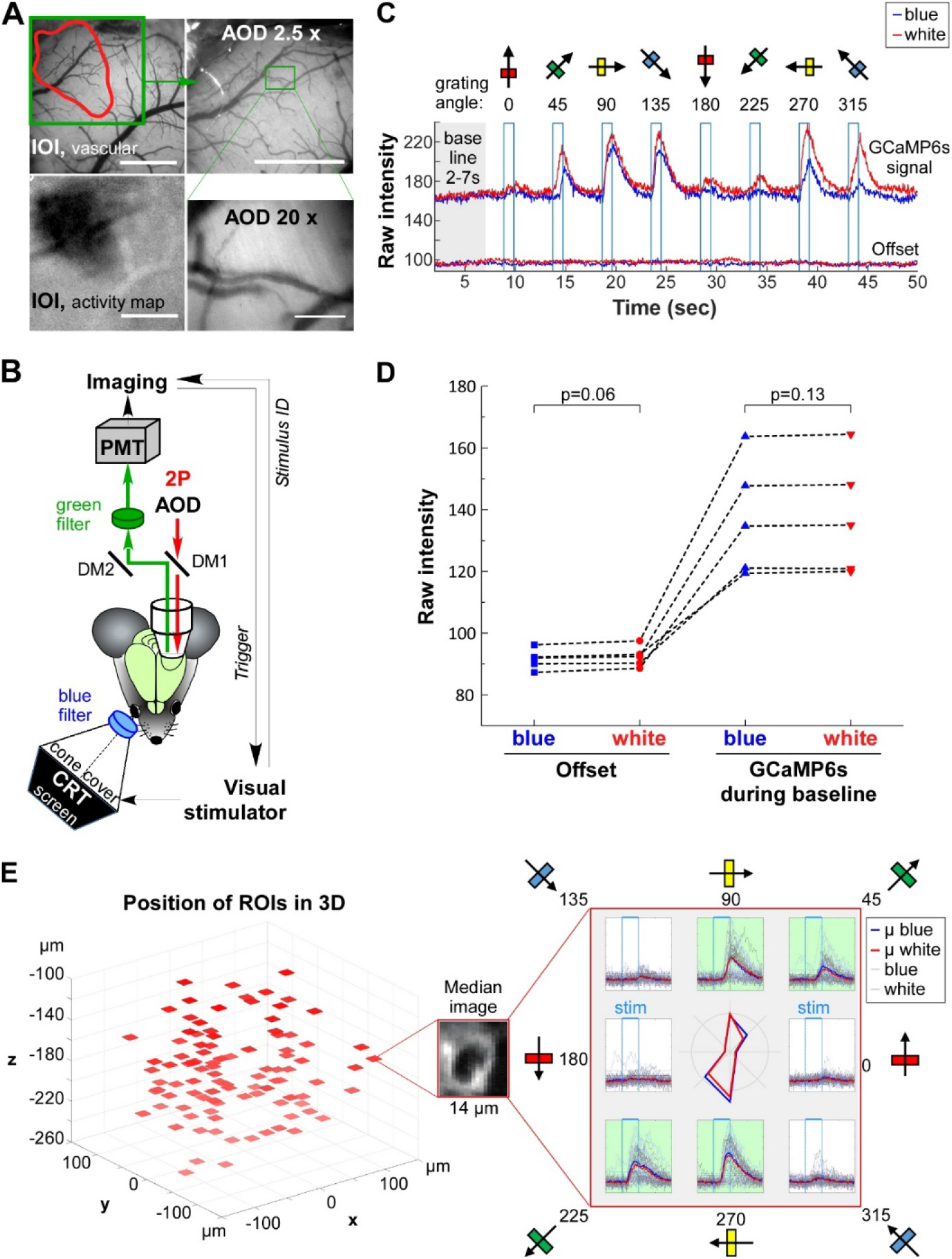
Experimental design and processing of Ca^2+^ imaging. **A**. Selection of the region for Ca^2+^ imaging. Example obtained by optical imager of vascular image (upper left) overlaid with outlined in red visual activity map obtained by IOI (lower left). The same area in camera mode under AOD setup at low magnification (2.5x; upper left) with the region selected for Ca^2+^ imaging (outlined in green). The same region at higher magnification (20x, lower right). The scale bar for IOI images and AOD 2.5x is 1mm and for AOD 20x – 100 µm. **B**. Sketch of 2P Ca^2+^ imaging using AOD setup, performed in V1 of Thy1-GCaMP6s mice during visual stimulation, presented on the CRT screen. In *italic*: triggering and timing control of the system. DM1 and DM2 dichroic mirrors 1 and 2. **C**. Example of time courses of (1, above) raw intensities of GCaMP6s fluorescence plotted for individual ROI during two single acquisition trials and (2, below) offset values in representative experiment – intensities averaged over all ROIs while laser shutter was closed. Intensities during blue/black (in blue) or white/black (in red) gratings presentation. Note that “blue” and “white” offset values overlap along entire time course; as well do the raw intensities of GCaMP6s fluorescence during the baseline period (2-7s; highlighted in grey) used for **F**_**0**_ calculation, suggesting comparable level of light contamination between two stimulation types. Stimulation periods (1s each) are outlined with blue rectangles; directions of drifting gratings are indicated above. **D**. Mean offset values (first 2 columns) and mean raw GCaMP6s fluorescence values during baseline period (**II** in C; last 2 columns) upon blue/black (in blue) or white/black (in red) gratings presentation. Values are averaged across all ROIs, all trials and time (of entire acquisition for offset or period within 2-7s for raw GCaMP6s fluorescence) within 1 experiment (1 animal). Squares, circles or triangles represent single animals (N=5). **E**. Left: example of acquired neurons located in 3D space. In the middle - median RIO image of representative cell, created across the data frames within the experiment. Right: responses (ΔF/F_0_) of this cell to blue/black (in blue) or white/black (mean – in read, single recordings - in gray) gratings plotted separately for each of 8 directions; thick lines: mean ΔF/F_0_ (µ) averaged over the trials (n = 21 and 17 for blue and white respectively); thin: ΔF/F_0_ of individual trials; in the middle – polar plots of the responses evoked by blue/black (in blue) or white/black (in red) gratings. Stim – time of the grating presentation.

After a pixel by pixel subtraction of the first 10 frames from subsequent ones, activity maps were obtained by dividing the images of cortex, stimulated with the gratings, by the average blank image of the unstimulated cortex (to correct for uneven illumination; (Grinvald et al., 1999)). The obtained maps of visual response to 0 or 90 degree gratings were largely overlapping (Supplementary Fig 2.1). The visually active part of V1 was found by overlaying the activity map with vasculature image acquired before (see Fig. 2A upper left, visually active area is outlined in red). The activity map was used to guide subsequent 2P Ca^2+^ imaging. The vascular pattern of the active area was used as a landmark for alignment during 2P set up using a low magnification objective (Zeiss 2.5x air objective, 0.06 NA; see Fig. 2A upper right) with help of the microscope camera system. Part of these area was chosen for Ca^2+^ imaging, performed under higher (20x) magnification (see Fig. 2A lower right).

#### 2.3.3. Ca^2+^ imaging

2P Ca^2+^ imaging of local populations was performed using a 2P microscope with acousto-optic deflectors (AOD, Femtonics femto 3D Omega), providing random access acquisition of multiple cell somata in 3D (Szalay et al., 2016). Data acquisition was controlled by the MES software (Femtonics; for setup sketch see Fig. 2B). The laser source was a Ti-Saphire laser (Tsunami 3941C-25XP femtosecond laser), running at 875 nm, pumped with Millennia eV 25S (Spectra-Physics). The objective used for Ca^2+^ imaging was a 20x water immersion lens with high numerical aperture (NA 1.0) and 2mm working distance (Olympus XLUMPLFLN-W). As an immersion fluid ultrasound gel Aquasonic 100 (Parker) was used. To minimize light contamination and the differences in offset value while presenting gratings of different colors, the gap between objective and the bath was masked with opaque plasticine. The power used to excite GCaMP6s was on average 84 mW (range: 73-106). The GCaMP6s expressing somata (with surrounding background areas) were selected from a 3D stack that was acquired before recording within the L2/3 (~80-260 µm under the pia). We used a chessboard scanning approach ((Szalay et al., 2016); see Fig. 2E, left) where for each chosen cell a 14×14 µm region of interest (ROI, with 0.7-0.8 µm pixel size) was acquired. The scanning speed was on average 21.24 Hz (range: 18.6-24.3; depending on the number of ROIs). To estimate amount of light contamination (offset) during given stimulation type, the acquisition was repeated with the laser shutter closed.

**Supplementary Figure 2.1.**
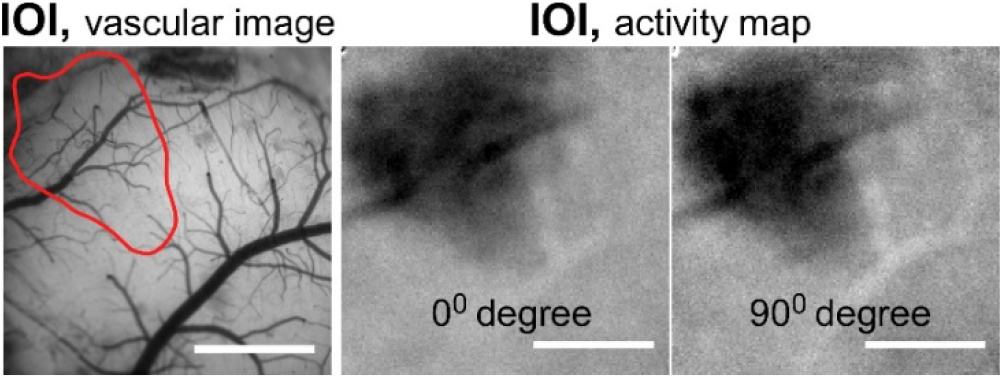
From left to right, respectively: vascular image (area detected to be active during visual stimulation is outlined in red), activity maps obtained during the presentation of gratings, drifting at 0^0^ or 90^0^. Note that activity maps are similar. Bar: 1mm

#### 2.3.4. Visual stimulation during Ca^2+^ imaging

As during LFP recordings, the CRT screen was shielded by an opaque cone with a slit for insertion of the blue filter and placed at 40 cm from mouse eye, centered at an angle of 40-45 degree from the vertical meridian. Visual stimuli were drifting blue/black or white/black gratings (with sine waveform, spatial frequency 0.03 cycles/degree, drift velocity 1 cycle/s). As it was described in the section *2.2.3* (“*Visual stimulation during LFP recording*”) white/black and blue/black gratings had equal output powers and the “blue” stimulus was achieved as 100% white on the CRT screen + blue filter (see Fig. 2B). During each stimulus trial, sinusoidal drifting gratings of 8 different directions (0, 45, 90, 135, 180, 225, 270, 315, 360 degree angle) were presented. Stimulation with each grating direction lasted 1s, with a minimum 3s of blank period before (see Fig. 2C). We used “compass” coordinates, in which horizontal gratings moving upward are considered to be moving at 0°, and angles increase in a clockwise manner (Mazurek et al., 2014). Somata were acquired for 50 s (one trial duration). Before, after stimulus trial and in-between gratings equiluminant blanks were presented. The acquisition trial was repeated at least 10 times for each stimulation type (white/black or blue/black gratings). After 5 s of spontaneous activity recording, the MES 2P data acquisition system triggered the whole visual stimulation cycle (8 differently oriented grating presentations + blanks). The time of grating presentation (managed by ViSaGe system) and the grating direction ID were always recorded in MES software (via analog input to align and associate the data frames with grating ID (see Figs. 2B,C)).

A higher background contamination during white/black gratings presentation due to white light component could have caused a higher offset, thereby higher baseline fluorescence of GCaMP6s (F_0_) and, consequently, spurious similarity of the calcium transients between two conditions, even in case of weaker response to blue/black gratings. We thus took the countermeasures and masked the gap between objective and the bath with opaque plasticine. We then recorded the signal from selected cellular ROIs while the laser shutter was closed. In these conditions, obtained signal (offset) should be only due to the contamination from ambient environment (e.g. from the CRT screen), dark noise and noise from the electronics. **Fig. 2 C** shows that the representative “offset” time courses recorded in these configuration during blue/black and white/black gratings presentation strongly overlapped. Averaged over the time of acquisition “offset” values were slightly bigger during white/black gratins presentation, but the difference did not reach statistical significance (“offset”: 91.55 ± 1.46 and 92.39 ± 1.52; p = 0.06 for blue and white gratings, respectively, Wilcoxon signed rank test; **Fig. 2 D**, first 2 columns). More importantly, the raw values of acquisition time course section, used for the “baseline” fluorescence (F_0_) calculation (2-7 s, during presentation of blank), averaged for all cells and trials within each experiment (referred to as “mean raw baseline”), were not significantly different during the presentation of blue and white blanks: 137.33 ± 8.36 and 137.66 ± 8.44, respectively; Wilcoxon signed rank test, p = 0.13; **Fig. 2 D**, last 2 columns).

#### 2.3.5. Analysis of Ca^2+^ imaging

Analyses were performed in MatLab. First, offset values (due to residual light contamination from ambient light and visual stimulation) were calculated for each experiment and stimulation type (blue/black or white/black gratings) as mean intensity over all pixels of each ROI over time course of acquisition trial with the laser shutter closed. This value was subtracted from all pixels from each ROI across entire acquisition duration. Later a circular standard mask was placed over the cytoplasmic region of the cell within each ROI. Masks’ positions were adjusted manually for each cell and each acquisition trial to correspond to the median image created across the data frames within that trial. The adjustment was meant to correct at least partially for brain motion artifacts. For each data frame, the intensities of the pixels within and outside the mask were averaged, providing the values **F**_**cell_measured (t)**_ and **F**_**neuropil (t)**_ respectively (for the cell and neuropil). The fluorescent signal of the cell body was calculated as:

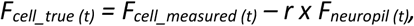

Although for optimal neuropil correction r should be adjusted depending on depth, we have chosen to use r = 0.7 across preparation in order to compare our data with those obtained with conventional mirror-based scanning systems with the same GECI (GCaMP6s) we had (either virally delivered or stably expressed in the same mouse strain we used here) and with a similar visual stimulation protocol, where the same r value was used for all neurons (Chen et al., 2013; Dana et al., 2014). Moreover, r has been reported to vary within a limited range within L2/3 (of ca 0.1, e.g. in (Kerlin et al., 2010)). The fluorescence time course of the cell body was plotted and aligned to the onset of individual grating presentation.

The baseline fluorescence (**F**_**0**_) was estimated as the intensity corresponding to 20% of the cumulative distribution function of F_cell_true (t)_ within 2^d^-7^th^ s of acquisition. **ΔF/F**_**0**_ was calculated as:

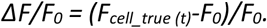

In order to transfer the ΔF/F_0_ time courses of each acquisition trial into polar coordinates, they were split into 8 segments, each for a single direction of drifting gratings with 1s of pre-stimulus, just before stimulus onset, and 3s after it (see Fig. 2E, right). For each direction the blank value was defined as ΔF/F_0_ averaged over 1s of pre-stimulus. Then, mean blank value across single “blue” or “white” acquisition trials segments (recorded during blue/black or white/black grating presentation, respectively) was calculated for a given cell and direction and subtracted from ΔF/F_0_ values of single trial segments to bring the pre-stimulus to zero. Single and mean “blue” and “white” ΔF/F_0_ (**µΔF/F**_**0**_) time courses were then plotted for each cell and direction (see Fig. 2E, right).

For each trial and direction, visual responses of individual cells were defined as the maximum ΔF/F_0_ within 2s, starting from stimulus onset. For a neuron to be qualified as visually responsive two conditions had to be simultaneously satisfied: (1) its visual responses for each orientation had to be significantly different from the corresponding blank value using a paired statistic (Wilcoxon signed rank test, p<0.01); (2) its **max µΔF/F**_**0**_ within 2s from the stimulus onset (referred to as **mean response** or **response amplitude**) had to be above mean blank (pre-stimulus µΔF/F_0_) + 2 SD.

The orientation (**OI**) and direction (**DI**) indexes were calculated for visually responsive cells (Mazurek et al., 2014). Preferred orientation (**R**_**pref_ori**_) was determined for blue/black or white/black gratings as orientation with largest magnitude of the response among the 4 orientations. Magnitude of each orientation was calculated as sum of max µΔF/F_0_ of two corresponding opposite directions. The orientation orthogonal to R_pref_ori_ (**R**_**orth_ori**_) was defined as *R*_*pref_ori*_ + 90°. OI was computed as:

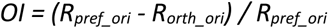

Preferred direction (**R**_**pref**_) for blue/black or white/black gratings was determined as direction with largest max µΔF/F_0_ among the 8 gratings directions presented. Null direction (**R**_**null**_) was defined as direction opposite to R_pref_ (*R*_*null*_ *= R*_*pref*_ + 180°). DI was calculated as:

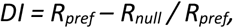

To quantify relative responsiveness of neurons towards blue/black versus white/black gratings, we defined a chromatic preference index (**cPI**, (Aihara et al., 2017; Tan et al., 2015)) as

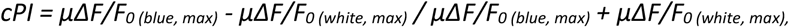

where **µΔF/F**_**0** (blue, max)_ and **µΔF/F**_**0 (**white, max)_ are the mean responses of an individual cell to preferred direction evoked by blue/black and white/black gratings, respectively. cPIs of 1 and −1 indicate complete blue/black and white/black dominances, respectively. Neurons giving identical responses to blue/black and white/black gratings have a preference index equal to “0”.

### 2.4. Statistical analysis

The statistical analysis was done in Origin 2017 and in MatLab. To compare two paired groups with low sample size or groups that were significantly different from normally distributed populations we used non-parametrical paired Wilcoxon signed-rank test. Normality was examined using Shapiro-Wilks test. To compare two paired normally-distributed groups paired t-test was used. Multiple matched groups (percentage of responsive to white, blue, or both gratings) were compared with the non-parametric Friedman test. Two independent groups (mean response of cells responsive to only blue or white gratings vs those of cells responsive to both) were compared with non-parametrical Mann-Whitney test. Cumulative distributions (of OIs and DIs) were compared by Kolmogorov–Smirnov test. The relationship between OI or DI of cell in response to blue and white gratings was estimated by linear fit function slope (**k**) and Pearson correlation coefficient (**r**).

## 3. Results

We first selected a “blue filter” transmitting a light spectrum(392-497 nm) from the CRT screen that would not overlap with the acquired GCAMP6s emission spectrum, band-passed by the “green filter” (503-539 nm, FBH520-40, Thorlabs) before the microscope’s photomultiplier tube (PMT) (see **Fig 1A and 2 B**). The “blue” filter was inserted into the slit at the tip of an opaque frustum CRT-screen cover (see **Fig 1B and 2 B**) in front of mouse eye. To compare the responses to blue and white, the intensities of the stimuli were adjusted to equal powers at eye level (ca. 3 µW): CRT intensity was set to 100% for blue and 35% for white (see section *2.2.3* - “*Visual stimulation during LFP recording*”).

### 3.1. Blue/black and white/black gratings trigger visual evoked potentials with similar amplitudes and peak latencies

To estimate the effectiveness of the blue light in driving mouse V1 activation compared to the white light, we first recorded VEPs in layer 2/3 in response to blue/black or white/black stationary gratings alternating in counter-phase presented to the contralateral eye of anesthetized mice, adapted to the ambient light level of the set-up used for physiology experiments (**Fig. 1 B;** for details see the section *2.2.3* - “*Visual stimulation during LFP recording*”). Blue/black gratings evoked similar responses compared to the ones evoked by white/black gratings, as shown by the largely overlapping VEP grand average traces recorded in the two conditions (N=8 mice; **Fig. 1 C**). A paired comparison of the peak amplitudes and latencies of VEPs, determined within 0.3 s from stimulus onset, showed that these parameters were not significantly different (blue vs white, peak amplitudes: −0.148 ± 0.051 vs −0.140 ± 0.047 mV; peak latencies: 0.235 ± 0.021 vs 0.230 ± 0.012 s; Wilcoxon signed rank tests, p=0.55 and paired t-test, p=0.83, respectively; **Fig. 1 D,E**). Thus local synaptic inputs into the V1 network in response to white and blue light stimulations, as represented by VEPs, had similar magnitudes and reached the network with comparable latencies.

### 3.2. Blue/black and white/black gratings activate two strongly overlapping neuronal subnetworks to a similar extent, as revealed by 2P population Ca^2+^ imaging

We next addressed the same question, at supra-threshold (spike output) level using 3D 2P population Ca^2+^ imaging in layer 2/3 of V1, performed with an AOD 2P microscope (see the section *2.3.3.* – “*Ca*^*2+*^ *imaging”*), technique which allows fast network monitoring with single cell-resolution. We compared the cellular response to blue/black and white/black gratings drifting in one of eight different directions presented to the contralateral eye of lightly-anesthetized Thy-1-GCaMP6s mice (see the section *2.3.* – *3D Acousto-optic 2-photon Ca*^*2+*^ *imaging of neuronal populations)*. See also entire **Figure 2**.

After defining the cells with a significant visual response to at least one direction of the gratings (see the section *2.3.5.* – “*Analysis of Ca*^*2+*^ *imaging”*), we calculated the percentage of the neurons responding to either stimuli in each experiment (N=5 mice; 81-100 neurons were recorded per animal, and 452 in total). Blue/black and white/black drifting gratings activated a similar percentage of V1 neurons (**Fig. 3 A**; p = 0.88, Wilcoxon signed rank test; on average: 13.9±3.6% and 13.5±3%, respectively). Besides, the fraction of cells, responsive to both color-types of gratings, was also comparable to abovementioned values (**Fig. 3 A**; Friedman test; p = 0.12; on average: 10.9±2.4%), suggesting that two populations – responsive for blue and for white - strongly overlap. Indeed, across all responsive cells (N=75) the largest portion (66.7%, 50 cells) responded to both color-types of gratings (**Fig. 3 B**), meaning that blue and white activated largely the same neuronal subnetwork. Smaller populations of cells were responsive only to the blue/black (18.7%, 14 cells) or only to the white/black (14.7%, 11 cells) gratings (**Fig. 3 B** and **Supplementary fig. 3.1**). A closer examination of these cells revealed that their mean responses (max µΔF/F_0_ values) in the preferred direction were lower than those of the cells responding to both color-types of gratings (in response to blue: 0.63±0.07 for cells responsive to blue only versus 1.07±0.10 for cells responsive to both, p = 0.03, Mann-Whitney test (**Supplementary fig. 3.1 C**); in response to white: 0.56±0.05 for cells responding to white only versus 1.05±0.09 for cells responding to both, p = 0.006, Mann-Whitney test (**Supplementary fig. 3.1 D**)).

**Figure 3.**
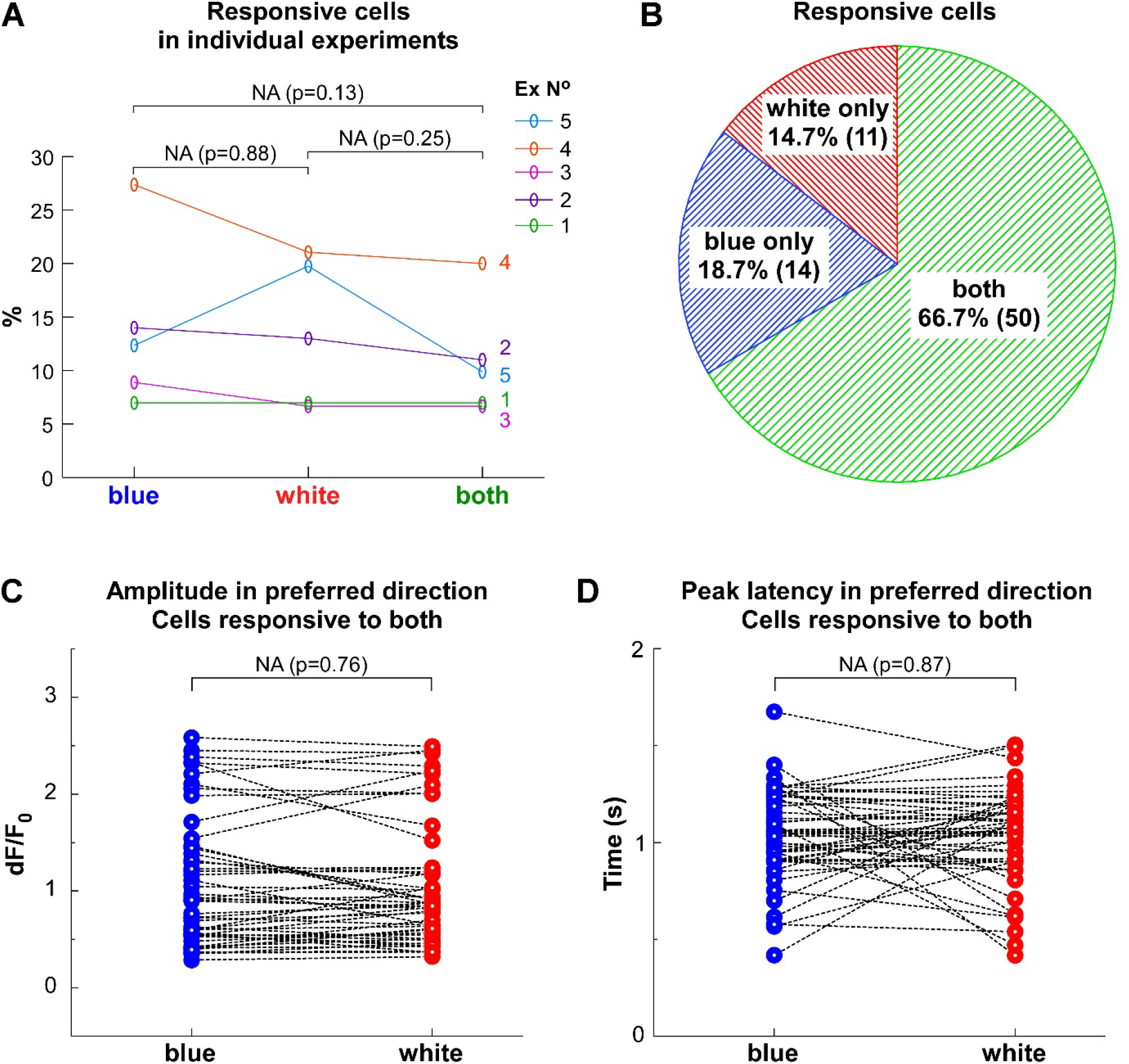
2P population Ca^2+^ imaging reveal that blue/black and white/black gratings activate strongly overlapping neuronal subnetworks to a similar extent. **A**. Percentage of cells in individual experiments (N=5) responding to blue/black, white/black and both gratings. Note, that there is no statistical difference in percentage of cells respond to blue/black versus white/black gratings (p=0.88; Wilcoxon signed rank test). **B**. Pie chart of cells responding to only blue/black or white/black gratings, or to both gratings (data pulled together from all experiments). **C, D**. Amplitude (**C**) and the peak latency (**D**) of mean Ca^2+^ responses in the preferred direction (in cells responsive to both blue/black and white/black gratings). Circles represent single cells. Values are paired for each cell. Note, that there is no statistical difference between the groups neither in amplitude nor in peak latency (p=0.76, Wilcoxon sign-rank test and p=0.87, paired t-test, respectively).

We next tested whether blue and white visual stimuli, beyond activating common V1 subnetwork, triggered similar responses. So, we analyzed the amplitude (= mean response = max µΔF/F_0_ values) and the peak latency of the visually-driven Ca^2+^ responses in cells responsive to both stimuli. Both response parameters upon stimulation with white/black vs blue/black gratings drifting in the cell’s preferred orientation were similar at the single-cell level (**Fig. 3 C, D**; n = 50 cells (from 5 mice); on average, peak amplitudes: 1.05±0.09 vs 1.07±0.10, and peak latencies: 1.03±0.03 vs 1.04±0.03 s; p=0.76, Wilcoxon signed rank tests; and p=0.87, paired t-test, respectively). The same was true in all the other (non-preferred) directions: the differences between response to blue and to white in their amplitude’s and latency’s values, calculated separately for each cell and each direction, were clearly distributed around zero (see **Supplementary Fig. 3.2**). Noticeably, the temporal profiles of Ca^2+^ transients visually-evoked by blue and white were strikingly similar (**Supplementary Fig. 3.3)**. Thus, the degree of visual responsiveness to white/black and blue/black gratings was similar for both the preferred and non-preferred orientations for major part V1 network responsive to both stimuli.

To estimate the color preference of the whole neuronal population, we looked at the distribution of the chromatic preference index (cPI; see the section *2.3.5.* – “*Analysis of Ca*^*2+*^ *imaging”*). cPIs distributions (1) of the cells responding to both blue and white gratings (N=50; **Supplementary Fig. 3.4 A**) as well as (2) of all responsive cells (N=75; **Supplementary Fig. 3.4 B**) were normally distributed (Shapiro-Wilks tests, p=0.41 and p=0.59, respectively) and had mean values approximating 0 (−0.002 ± 0.018 and −0.003 ± 0.02, respectively), suggesting that on the level of neuronal population there was no color preference.

**Supplementary fig 3.1.**
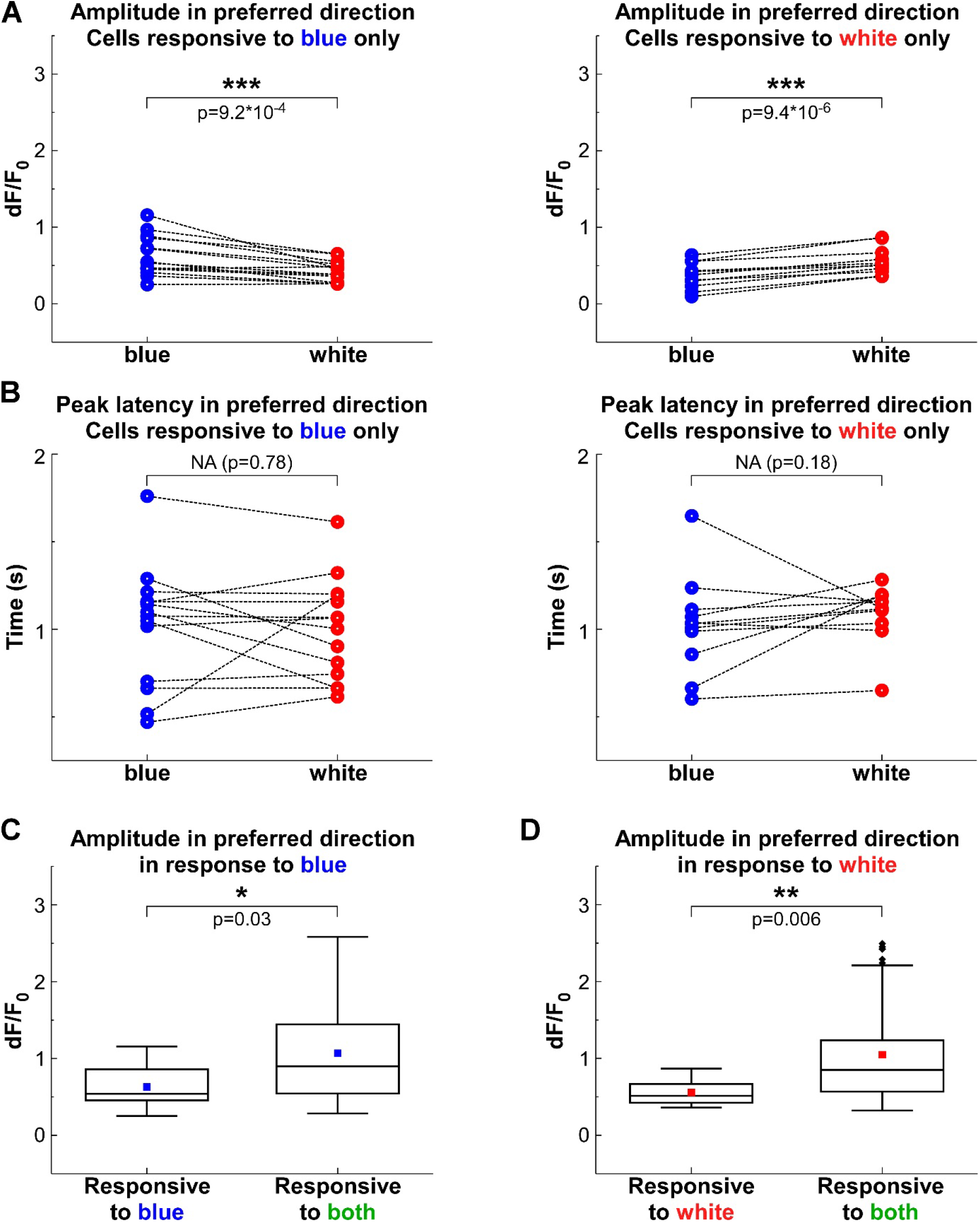
Amplitude (**A**) and the peak latency (**B**) of Ca^2+^ response in the preferred direction in single cells responding to only blue/black (on the left) or white/black (on the right) gratings. **C, D**. Comparison of the responsiveness to blue/black (C) or to white/black (D) gratings in cells responsive to only one color-type of grating (left column, to only blue in C or white in D) versus cells responsive to both stimuli (right column). Blue/red dots represent mean, box −25-75% of distribution with the median in the middle, whiskers – 1.5 interquartile range (IQR) and black dots – outliers.

**Supplementary Fig 3.2.**
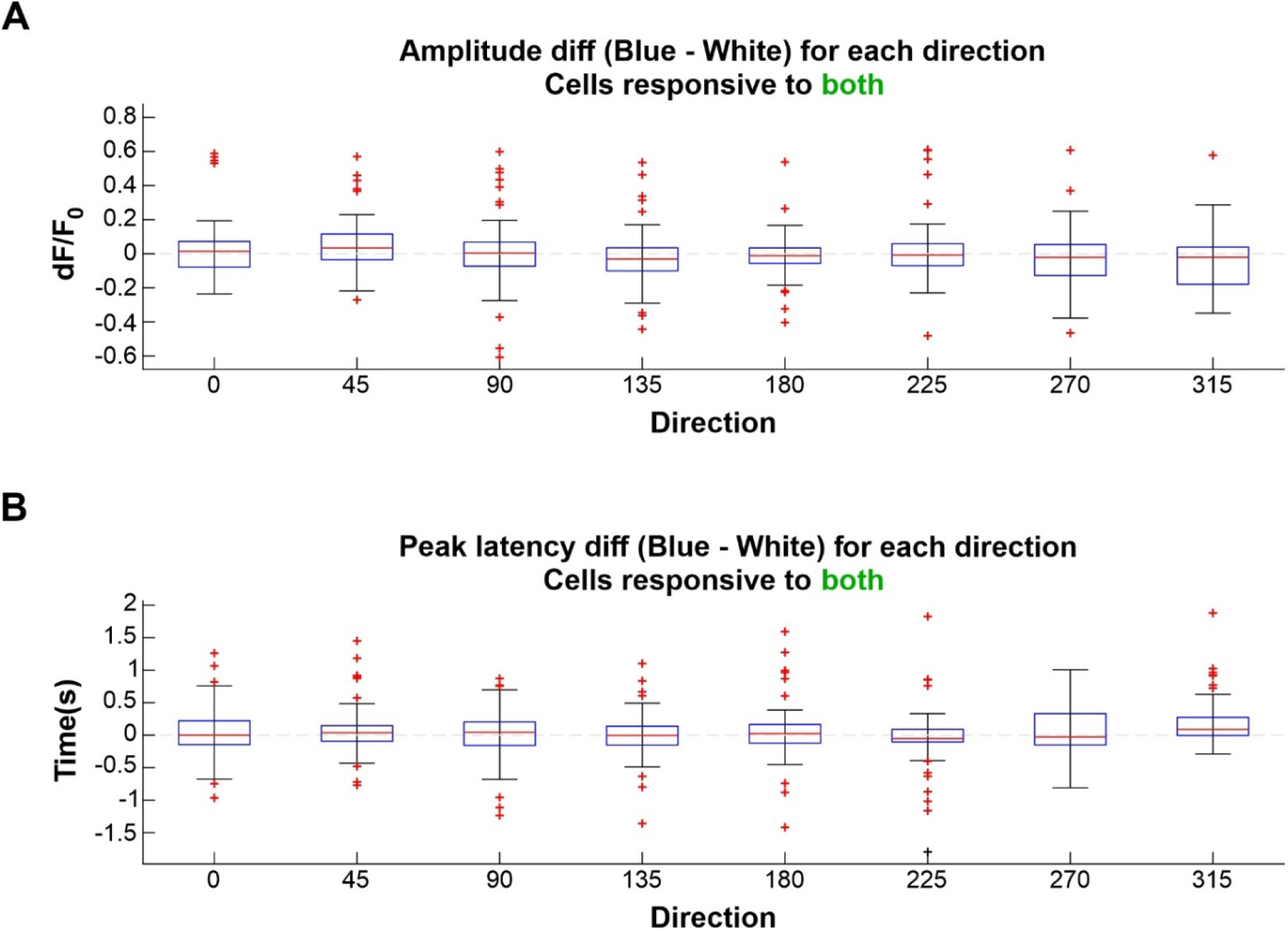
Differences between blue-and white-driven Ca^2+^ responses for each direction. Amplitude (A) and the peak latency (B) of Ca^2+^ responses. The differences were calculated for each cell responding to both gratings. Red line - median of the difference across the cells; blue box −25-75% of distribution; bars: extreme data points; red crosses: outliers. Horizontal gray dashed line is equal to 0.

### 3.3. Orientation and direction selectivities are comparable upon blue/black and white/black gratings presentation

Taking into account that direction and orientation selectivities are important functional response properties of V1 neurons, we next examined whether these parameters are different upon blue and white visual stimuli presentation. Polar plots obtained from the responses evoked by blue/black or white/black gratings in cells activated by both stimuli looked very similar (**Fig. 4 A, B**). Polar plots of individual cells with robust responses to these two colors (with mean Ca^2+^ responses in preferred direction above the median response of the population of responsive to both stimuli cells) were overlapping as well (**Fig. 4 C)**. In line with that, the majority of the cells activated by both colors shared the same preferred orientation and direction in response to blue and to white, as the difference in these variables in degrees, between the two colors was close to zero (**Supplementary fig 4.1**).

**Fig 4.**
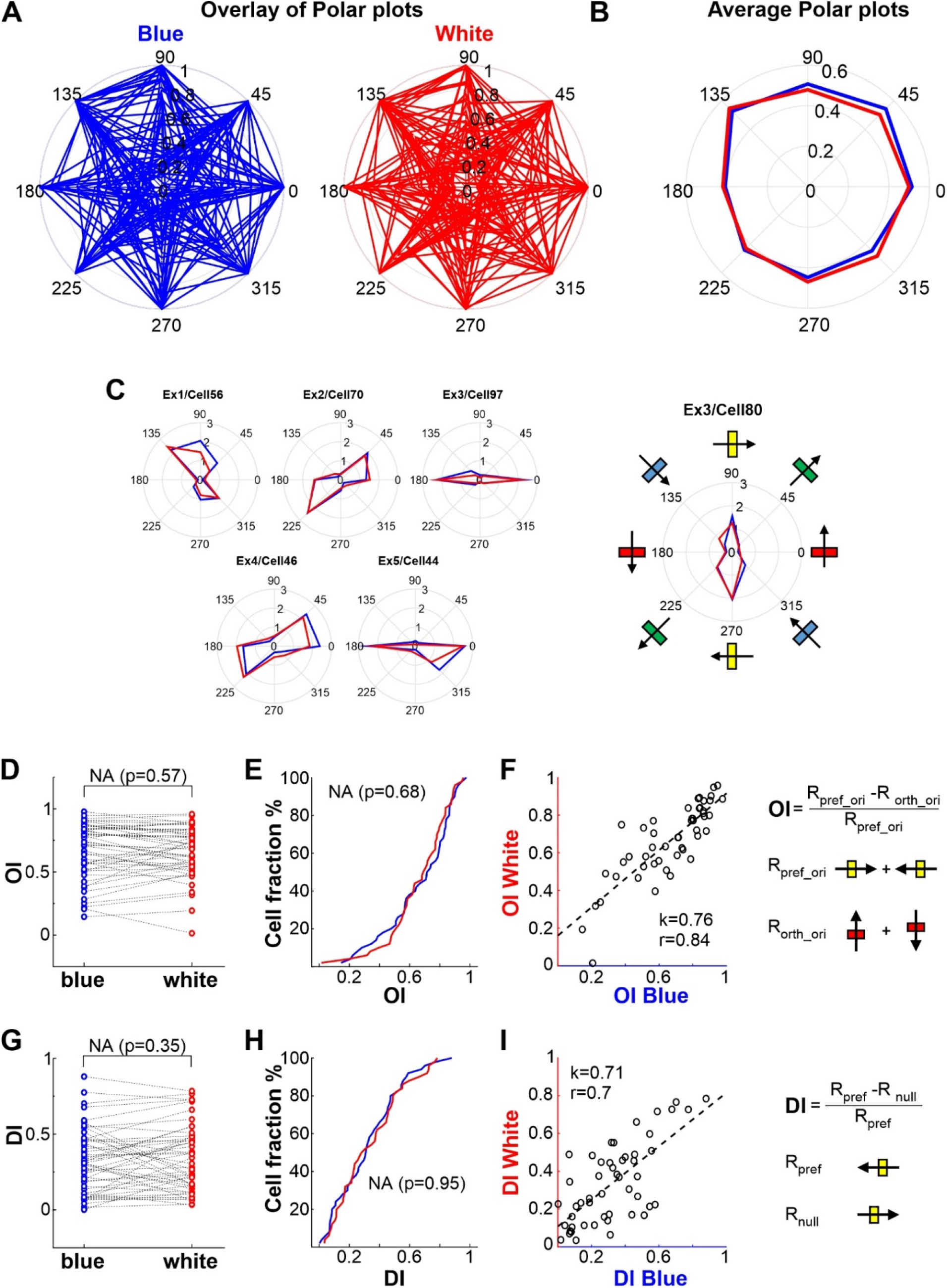
Orientation and direction selectivity of cells are comparable upon stimulation with blue/black and white/black gratings. **A**. Overlay of polar plots obtained from the responses evoked by blue/black (left) or white/black (right) gratings in individual cells. Mean Ca^2+^ responses for each direction are normalized to the maximal response (to preferred direction). **B**. Mean polar plots obtained from plots in A. **C**. Examples of overlaid polar plots for robustly responding cells (with mean Ca^2+^ responses in preferred direction above the median response across the population of responsive to both stimuli cells). Note that the polar plots are strongly overlapping. **D, G**. OI_s_ (**D**) and DI_s_ (**G**) of single cells in response to blue/black and white/black gratings. Values are paired for each cell. Note, that there is no statistically significant difference between the groups neither in OI nor in DI (p=0.57 and p=0.35, respectively, Wilcoxon sign-rank test). **E, H**. Cumulative distribution of OI (**E**) and DI (**H**). There is no statistical difference neither in OI nor in DI of responses evoked by blue/black and white/black gratings presentation (p=0.68 and p=0.95, respectively, Kolmogorov–Smirnov test). **F, I**. Scatter plots of OI (**F**) and DI (**I**) of single cells in response to blue/black (x-axis) and to white/black (y-axis) gratings. Circles represent a cell. Slopes of fitting linear functions (0.76 and 0.71 for OI and DI, respectively) suggest strong positive relationship between blue-and white-OIs/Dis; and Person’s coefficients (r = 0.84 and 0.7, respectively) - a strong correlation between them. Properties of response to blue/black gratings are in blue, to white/black – in red. All plots are for neurons that respond to both gratings.

We further looked at the degree of preference (described by selectivity indexes: OI / DI). A paired analysis of the orientation indexes (OIs) of individual cells did not reveal any significant difference (**Fig. 4 D;** mean OIs for blue and white: 0.67 ± 0.03 vs 0.66 ± 0.03, respectively; Wilcoxon signed rank test; p=0.57). In line with that, we failed to find any statistically-significant difference between the cumulative distributions of OI in response to these two color-types of gratings (**Fig. 4E**; Kolmogorov– Smirnov test, p=0.68).

**Supplementary Fig. 3.3.**
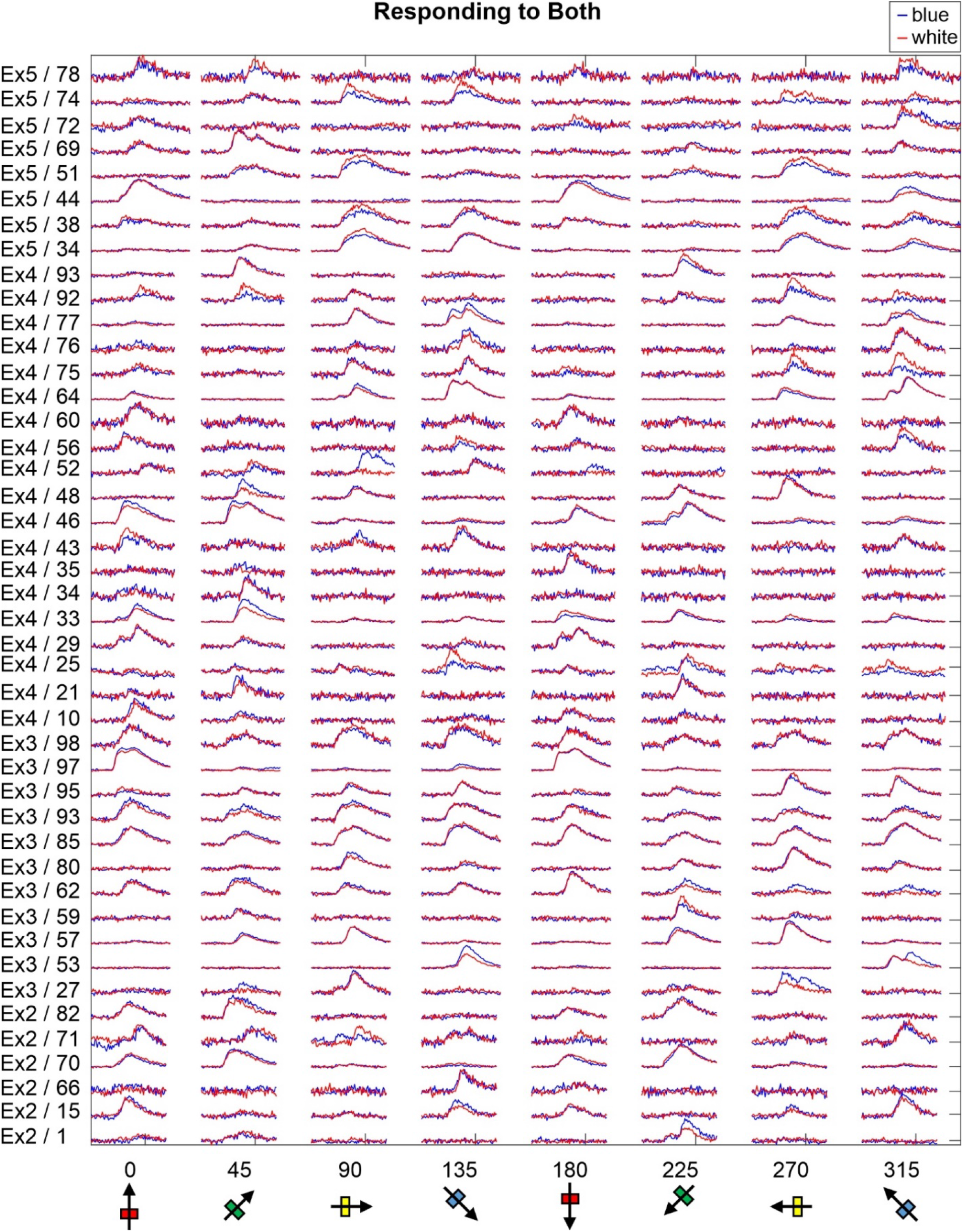
Time course examples of responses in neurons activated by both blue and white gratings. Responses are plotted for each direction (indicated below) separately and normalized to the maximal response for the cell. In blue and red – the responses (mean dF/F_0_) to blue and white stimuli, respectively. Note, that the shapes of mean Ca^2+^ transients in response to blue and white gratings for majority of the cells are strongly overlapping. On the left: the experiments / cell numbers.

**Supplementary Fig 3.4.**
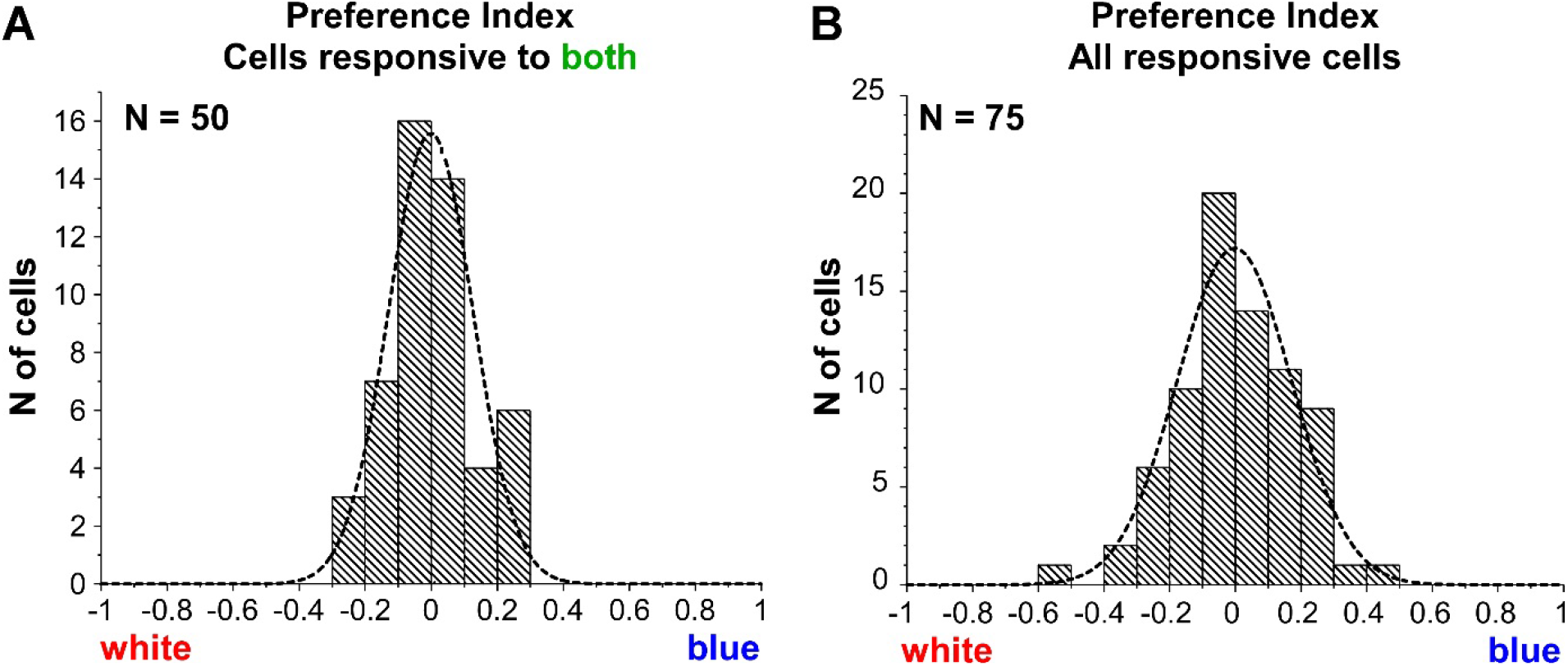
cPI distributions. Plots for cells responding to both gratings (**A**) and for all responsive cells (**B**). Note, that the distributions of cPI are normal (Shapiro-Wilks test, p=0.41 and p=0.59) with mean close to 0.

On the contrary we observed strong positive relationship between OI of cell in response to blue/black and OI in response to white/black gratings (linear fit function slope, k=0.76), in complement to strong correlation (r=0.84) (**Fig. 4 F**).

The directional selectivity of responses visually-evoked by white/black and blue/black stimuli was also similar. Indeed, we found neither difference in the paired analysis of the direction indexes (DIs) in response to white and blue color-types of gratings (**Fig. 4 G**; mean DIs: 0.34±0.03 vs 0.32±0.03, respectively; Wilcoxon signed rank test, p=0.35), nor in the cumulative distributions of this parameter (**Fig. 4 H**, Kolmogorov–Smirnov test, p=0.95) but instead we observed a positive relationship (k=0.71), as well as correlation (r=0.7) between them (**Fig. 4 I**). Of note, direction selectivity was less pronounced than orientation selectivity (mean OI = 0.67 and 0.66; while mean DI = 0.32 and 0.34 for blue and white gratings, respectively), in agreement with previous reports (Niell and Stryker, 2008).

**Supplementary Fig 4.1.**
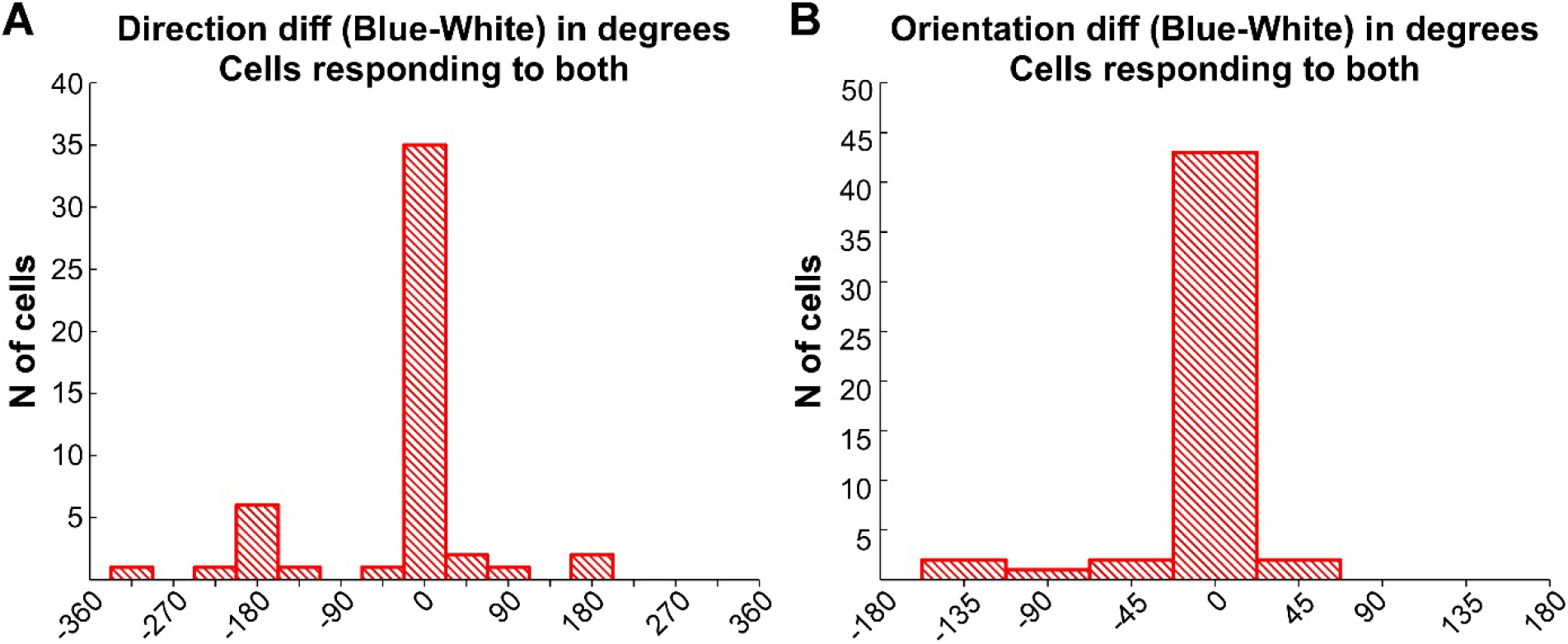
Histogram of difference in preferred direction (A) or orientation (B) of cells responsive to both gratings. Differences in degrees. Note that for majority of the cells this value is equal to “0”.

## 4. Discussion

White light (~ 380-730 nm) causes signal contamination in imaging experiments. Here, to minimize such contamination during Ca^2+^ imaging, we used blue light (~ 392-497 nm) instead, spectrally distinct from the acquired “green” GCAMP6s signal (as filtered before PMTs: 503-539 nm; **Fig. 1 A**). We might have expected that blue stimulation should have evoked weaker visual responses, since blue light comprises a smaller portion of visible spectrum absorption of both the rods’ rhodopsin (peak: 498 nm) and cones’ “green” (M/L) opsin (peak: 508 nm), than the white does.

Our data suggests instead that stimulation with blue light results in a similar local synaptic inputs (as assessed by LFP) and triggers a comparable spike output in the same neuronal subnetwork (measured optically by random access 3D two-photon calcium imaging). Of relevance, fundamental functional response properties such as orientation and direction selectivity of V1 neurons were also similar.

### 4.1. Explanation of the comparable physiological response in V1 L2/3 to blue and white visual stimuli

With electrophysiological approach (LFP) we were able to show that blue and white visual stimuli evoked similar local synaptic inputs into L3/2 of V1. With Ca^2+^ imaging we could observe that blue and white activated to the same extent two largely overlapping subsets of excitatory neurons in L2/3. In addition, blue and white triggered responses displayed similar direction and orientation selectivity - hallmark of mammalian V1 (Huberman and Niell, 2011), especially pronounced for excitatory cells in L2/3 and L4 (Hübener, 2003; Niell and Stryker, 2008). Such a comparison of V1 responses to equi-powerful stimuli with partially overlapping spectra (white embraced the entire spectra of blue) has not been done before. However, comparison of V1 responses to two spectrally distinct colors, that match two main picks of mice spectral sensitivity to UV and green lights, has been performed both by electrophysiology (Ekesten and Gouras, 2008) and Ca^2+^ imaging (Aihara et al., 2017; Tan et al., 2015). These studies illustrated that the majority of neurons responded to both and were similarly orientation-tuned also to both colors, especially the ones with higher orientation selectivity (Tan et al., 2015).

*The similarity of neuronal V1 responses might be due to the fact that the two stimuli activated the same retinal photoreceptors to a similar exten*t. Indeed, rods and cones are the main input of the classical (image-forming) visual pathway relaying information to V1 and are often the limiting factor for visual perception. Indeed, knock-in mice expressing a human long-wavelength-sensitive (L) cone photopigment, alongside the usual pigments, can discriminate green from red (Huberman and Niell, 2011; Jacobs et al., 2007), an ability that mice lack otherwise (Jacobs et al., 2007; Jacobs et al., 2004). Additionally, wild-type mice can distinguish UV and visible radiation, exploiting the separated chromatic sensitivities of cone UV and green opsins (Jacobs et al., 2004). The integrated retinal response reflects the summation of native photopigments’ contribution (Rocha et al., 2016), and behavioural spectral sensitivity mirrors the retinal one (Jacobs et al., 1991; Jacobs et al., 2007; Jacobs et al., 2004). Finally, the chromatic tuning of V1 neurons depends on retinotopy and reflects the spatial distribution of retinal photopigments (Aihara et al., 2017; Ekesten and Gouras, 2008; Rhim et al., 2017).

*What could have been the relative contribution of rods and cones?* In other words, is the similar drive of V1 neurons observed for blue and white stimuli valid in scotopic or photopic conditions? We estimate that the visual responses we recorded were at least partly attributable to rod activation. Rods (97% of photoreceptors; (Peirson et al., 2018)), are very light-sensitive, but able to adapt to light conditions traditionally considered to be photopic in rodents (Tikidji-Hamburyan et al., 2017). So at the light powers used (3 µW at eye surface, corresponding to ~10^2^ cd/m^2^ or to 10^10^ photons/s mm^2^), rod photo-transduction should happen at saturating levels (Naarendorp et al., 2010; Umino et al., 2008). This could contribute to the comparable responses to white and blue stimuli, even though a smaller fraction of the “green” peak part of rhodopsin’s spectral sensitivity function is covered by blue radiation.

Cones are very sparse (~3% of photoreceptors; (Peirson et al., 2018)) and orders of magnitude less sensitive than rods (Naarendorp et al., 2010), but in mice they might take over vision at lower luminances than in humans (Naarendorp et al., 2010). Both blue and white visual stimuli used can activate cone’s “green” sensitive M/L-opsin (expressed in 74% of all cones; (Ortín-Martínez et al., 2014)), whose spectral sensitivity is very similar to that of rhodopsin. Since cones saturates above ~0.1 cd/m^2^ (Umino et al., 2008), we estimate that white and blue stimuli activated cones to a similar extent.

*Although different, the blue and white spectra were “functionally” similar in their capability of recruiting the integrated retinal responses*, because: 1) the red component (present in the white but not in the blue stimuli) can recruit the mouse retina only at much higher light intensities (Peirson et al., 2018); 2) mice do not possess cones with blue photopigment, so, a stronger blue λ component in the blue light than that in white should not activate any cones better. Photosensitive retinal ganglion cells (pRGC), which express melanopsin with blue peak sensitivity (~478 nm), are not of our concern, since they are not implicated in image-forming, V1 responses (Peirson et al., 2018).

### 4.2. Impact of similar V1 responsiveness to white and blue stimuli on behaviour

*Can mice distinguish behaviourally white from blue?* Mice can discriminate colors (Denman et al., 2018; Jacobs et al., 2004), but only in certain conditions (e.g. if stimuli are above horizon and if chromatic changes are rather large). However, animals were tested to discriminate distinct colours within a narrow non-overlapping spectra (Jacobs et al., 2004), or the change in UV-green hue (importantly UV and green hae non-overlapping spectra’s, that match peak sensitivities of UV and green opsins (Denman et al., 2018). Behavioural comparisons of chromatic discrimination capacity between two overlapping spectra are missing.

The color-discrimination ability is constrained by the spectral sensitivity differences of photopigments (cone’s UV and green opsins, and rod’s rhodopsin) and determined by chromatic-opponent cells responding at “ON” to one light spectra and at “OFF” to another. Surprisingly, not only cones, but also rods can be implicated in color opponency (Joesch and Meister, 2016). UV-green opponent cells were found in some RGC (12% (Ekesten and Gouras, 2005; Ekesten et al., 2000)), and in V1 (~ 1% (Ekesten and Gouras, 2008; Tan et al., 2015)). We do know whether white-blue opponent cells exist in mice V1. It is difficult to extrapolate this information from the above mentioned studies, since the latter compared non-overlapping colors. However, since white and blue spectra overlap and do not engage different types of photopigments, the existence of opponent cells is questionable.

Apart from chromatic opponency, which has to do with the receptive field spatial substructure, neurons selectively responding to certain color or responding substantially stronger to one (here, with cPI considerably different from “0”) might also contribute to color-discrimination ability. Color-selective cells in L2/3 of mice V1 were also shown before (Aihara et al., 2017; Ekesten and Gouras, 2008; Tan et al., 2015). Using our criteria for responsiveness (see *2*.*3*.*5*.), we found neurons activated only by white or by blue. However, as we described in section 3.2., their responses were on average smaller than in cells responding to both colors and in some cells rather sparse. Thus, scarcer responsiveness might have caused a spurious classification as “unichromatic” in some of these cells.

*Up to which degree can these results be generalized to other mice strains*? Our experiments were performed in pigmented mice. Albino mice have relatively fewer total cones (Ortín-Martínez et al., 2014; Whitney et al., 2011), fewer cones expressing green opsin and more UV-opsin-expressing cones with less-pronounced dorso-ventral gradient of the last (Ortín-Martínez et al., 2014), and fewer rods (Donatien and Jeffery, 2002). However, the retinal selectivity of the pigments is similar, thus suggesting that our findings could be extended to albino strains.

*Our visual stimuli do not significantly recruit the UV sensitivity of the mouse retina*, as they did not effectively stimulate UV-opsin and the UV parts of rhodopsin and green opsin’s absorption spectra. Adding UV radiation to blue stimuli would engage UV mice spectral sensitivity, without causing light contamination during Ca^2+^ imaging. CRT monitors lack UV-spectra, but recently designed stimulators can do it (Franke et al., 2019), although this can raise concerns with respect to the lab safety.

*A limitation of our study is that we compared visual responses to blue and white stimuli in saturating conditions, that is with gratings of optimal spatial frequencies, temporal frequencies and contrasts*. Spatial and temporal frequency tuning curves, as well as a contrast sensitivity curve should be tested in order to have a more robust physiological basis for an educated guess on the behavioral equivalence of the two types of chromatic stimuli.

### 4.3. Generalized validity of results in different illumination conditions and different imaging techniques

*Both rods and cones should have contributed to the visual responses we recorded*. We performed experiments in conditions commonly used for *in vivo* electrophysiology and 2P microscopy: low level of ambient light and moderate intensity of visual stimuli (here 3 µW = 0.955 10^−2^W/m^2^ or ~10^2^ cd/m^2^, or ~10^10^ photons/s mm^2^, or < 1 lx) at eye level. This corresponds to mice photopic range (Umino et al., 2008). However, in these conditions rods continue to respond (~3 10^4^ R*/rod s (photoisomerization per rod) (Naarendorp et al., 2010) have been measured for luminances comparable to our conditions 10^2^-10^3^ cd/m^2^ (Umino et al., 2008)), even though they might temporarily loose sensitivity after abrupt luminance changes. In steady illumination, as in our experiments, rod’s contrast sensitivity, initially reduced in photopic range, was shown to recover (Tikidji-Hamburyan et al., 2017).

*We expect similar responsiveness in photopic and high mesopic* (above ~0.1 cd/m^2^ (Umino et al., 2008)) *conditions*. Indeed, both cones (Umino et al., 2008) and rods (Naarendorp et al., 2010) should respond at saturating levels in the luminance conditions used. The fact that our illumination level is approaching saturation is indeed suggested by the fact that a 3-fold light intensity increase (from our white 35% to white 100% intensity) caused on average only ~53% (much lower than 300%) increase in VEP amplitude (−0.15±0.05 vs −0.23±0.09 mV for 35% and 100%, respectively - see **Supplementary Fig. 1**). This contradicts older reports, showing quasi-linear amplitude increases upon luminance increment (Porciatti et al., 1999). On another hand, our finding is in line with recent behavioral studies. In one, carried out in mesopic conditions, authors documented that mice ability to discriminate two subsequent stimuli reached plateau already when the luminance difference between stimuli was just above 30% (Denman et al., 2018). In another work, mice contrast sensitivity plateaued at ~10^−2^ cd/m^2^ (Umino et al., 2008) – a luminance corresponding to our 35% white stimulus.

*We might expect possible differences in white-driven and blue-driven responses at low-scotopic conditions*. In such light conditions rhodopsin peak absorption should be critical, so the white light (embracing the entire rhodopsin peak sensitivity) might trigger larger responses than the blue light (that covers only half of it). Moreover, light spectrum can influence pupil constriction (Lucas et al., 2001), and the difference in pupil size upon blue and white stimuli can be noticeable in low scotopic conditions, but small in our luminance range (with pupil constriction ~ 85-90%).

*Blue visual stimulation, beyond allowing to omit time-consuming shielding, can facilitate to combine Ca*^*2+*^ *imaging with optogenetics* - so-called all-optical approach (Carrillo-Reid et al., 2017; Forli et al., 2018; Yang et al., 2018), *or with electrophysiological recordings* - e.g. (Chen et al., 2013). Electrophysiology requires indeed free access for recording electrodes, also rendering shielding problematic when combined with 2P experiments. All-optical microcircuit interrogation becomes even more problematic in head-fixed behavioral setups requiring rather “open” spatial configurations.

**Supplementary Fig 1.**
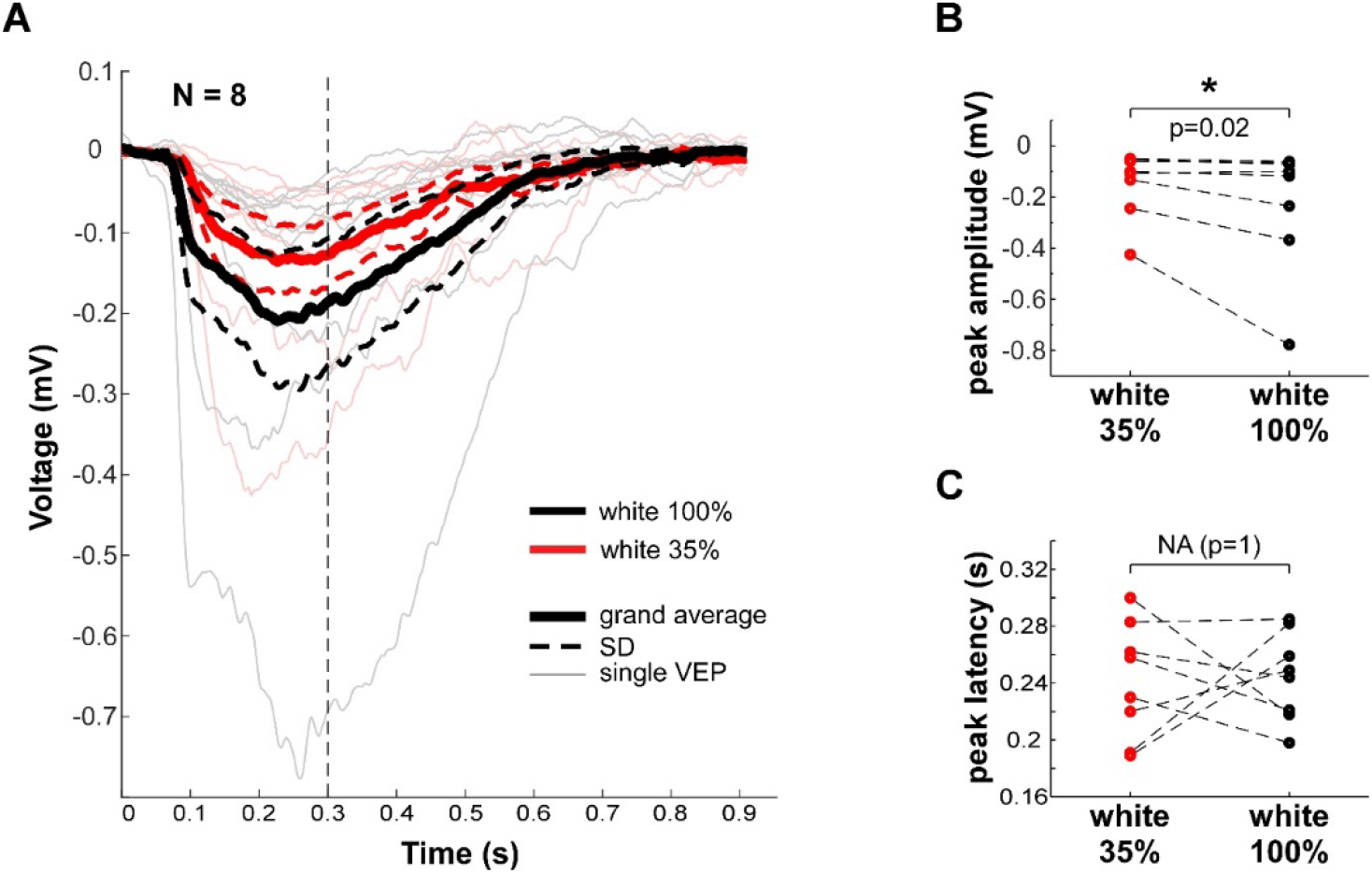
VEP in response to white/black gratings with 100% and 35% intensity of white in mice V1. **A**. Grand averages (N=8 mice, thick lines) ± SD (thinner dashed lines) and individual VEPs (thin lines) in response to 100% (in black) and 35% (in read) intensity of white. Dashed vertical line confines the time for peak search (0.3 s). Note that the responses are strongly overlapping. **B, C**. Amplitude (**B**) and latency (**C**) of single VEP’s peaks in response to 100% (in black) and 35% (in read) white intensity. Note, that VEP amplitude in response to 100% intensity is significantly (Wilcoxon signed rank test, p=0.02, N=8), but not 3 times higher than that in response to 35% intensity.

In relation to white, blue light stimulation can be beneficial while imaging not only GCaMP6s or other “green” cpEGFP-based sensors (λem = 515 nm) GECI (Chen et al., 2012; Chen et al., 2013; Dana et al., 2014; Dana et al., 2019), *but also red-shifted GECI* (as - jRGECO1a, jRCaMP1a and b; λem ~ 600 nm, (Dana et al., 2016; Dana et al., 2018)), since white (380-730 nm) visual stimuli would contaminate the acquired emission of the red-shifted dyes as well. *The application can be extended also to other sensors used to image neuronal function* such as synaptic activity reportrs, e.g. SynaptopHluorin (an indicator of synaptic release), or SuperGluSnFR and iGluSnFR (glutamate sensors) or voltage-sensitive dyes (Broussard et al., 2014; Carandini et al., 2015; Chemla and Chavane, 2010).

Despite utility of blue light, long-lasting high intensity blue visual stimulation should be avoided, as UV-violet-blue components of fluorescent lamps (with 4.97 10^−05^ mW/cm^2^/s radiant exposure) cause light-induced retinal damage within 1h of exposure (Narimatsu et al., 2014).

### 4.4. Comparison with alternative approaches

In addition to shielding, light contamination can be minimized by temporally separating the recording of fluorescence of Ca^2+^ indicator and visual stimulus presentation (Leinweber et al., 2014; Reiff et al., 2010), exploiting also the relative sluggishness of calcium responses compared to the voltage response. Stimuli are shown during the time periods when the data is not acquired/eliminated (during the turnaround points in case of raster scanning in the 2P scope). This, however, implies precise synchronisation of the screen’s/projector’s light output to the scanning/data collection of the imaging system and requires tailor-made electronics to precisely synchronize stimulation-acquisition times. In addition, this approach can impose serious constraints in the temporal structure of the presented stimuli.

## 5. Conclusions

Blue light, spectrally selected so not to interfere with calcium imaging with green-shifted GCAMP6s’ emissions, activates mouse V1 to a similar extent as an equi-powerful white light. This was assessed by measurements of both integrated synaptic responses (VEPs, reflecting the input) and Ca^2+^ responses of single neurons in the network (mirroring their action potential output (Kerr et al., 2005)). White and blue stimuli recruit the same network of neurons to a similar extent, without differences in fundamental functional response properties such as orientation and direction selectivities.

## Author contributions

PM supervised the project; PM and TK did experimental planning; TK performed the experiments and wrote the manuscript; PM, KA, EM, SP – revised the manuscript, KA – developed the Matlab code for VEP and Ca^2+^ imaging analysis, analyzed Ca^2+^ imaging in Matlab, developed visual simulation software and software communication for LFP, IOI and AOD-experiments; TK analyzed the data in Origin; KA and TK prepared the graphs; TK prepared final illustrations; EM and KA developed stimulation protocol for AOD and IOI; EM optimized the surgery and experimental design for Ca^2+^ imaging and IOI and preliminary experiments; SP contributed to Ca^2+^ imaging analysis.

Acknowledgements

Supported by grant Knut och Alice Wallenberg KAW 2014.0051 and Swedish Research Council (Vetenskapsrådet) VR 2014.02350.

We thank Sebastian Sulis Sato for the help with mastering the AOD set up, experimental and critical suggestions and Thy-1-GCaMP6s colony management; Sebastian Sulis Sato and Anushree Tripathi for showing the IOI; Johan Zakrisson and Sebastian Sulis Sato for tuning Ti-Saphire laser in AOD set up; Johan Zakrisson for cone slit design and light power measurements; Johan Zakrisson and Luciano Censoni for support in photometry units transfers; Per Utsi for help with mechanical engineering.

## Supplementary data

By visual examination we found 3 clear “off cells”, activated solely by the termination of the grating stimulation (with activation onset after stimulus OFF). Interestingly, the offset of both blue and white grating stimulation could trigger the increase in GCaMP6s fluorescence in these cells (see Supplementary Fig 2.2). As the time windows to calculate the response and blank values in our analysis were adapted for “on-cells”, “off cells” were not determined as responsive and not included in further analysis.

**Supplementary Figure 2.2.**
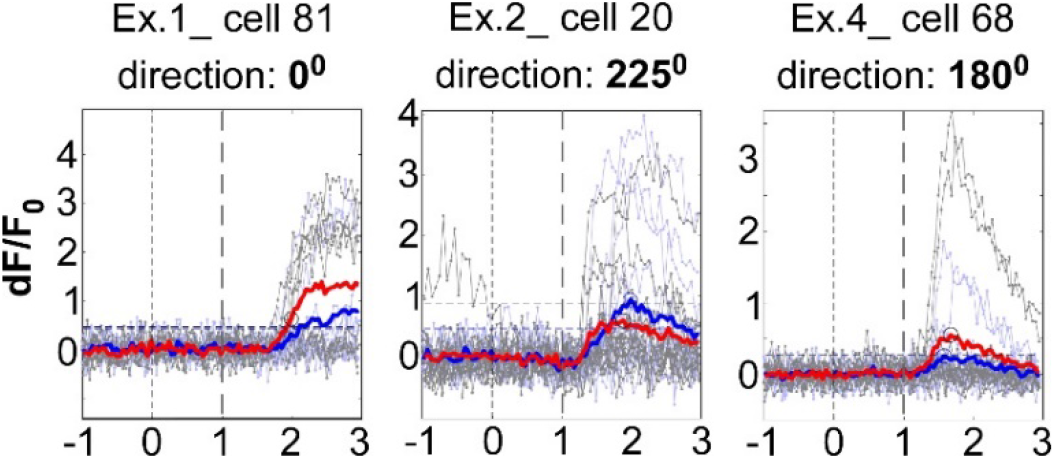
“Off cells”. Response of the given cells to one of the grating’s direction, indicated above. Thick lines: mean dF/F_0_ averaged over the trials (to blue and whitein blue and red, respectively); thin: dF/F_0_ of individual trials (to blue and white - in blue and grey, respectively). Above: experiment and cell numbers. Note, that the increase in GCaMP6s fluorescence is triggered by the termination of both blue/black and white/black gratings.

## Abbreviations

3D: three-dimensional
2P: two-photon
V1: primary visual cortex
VEP: visually-evoked potential
LFP: Local field potential
i.m.: intramuscular (injection)
AOD: acousto-optic deflectors
PMT: photomultiplier tube
ROI: region of interest
SD: standard deviation
cPI: chromatic preference index
OI: orientation index
DI: direction index
GECIs: genetically-encoded calcium indicators
EGFP: enhanced green fluorescent protein
CRT: cathode-ray tube
IOI: Intrinsic optical imaging
ERG: electroretinogramm

